# Non-redundant odorant detection in a locust

**DOI:** 10.1101/2022.06.21.496967

**Authors:** Hetan Chang, Anjana Unni, Megha Treesa Tom, Lucas Cortés Llorca, Sabine Brase, Sascha Bucks, Kerstin Weniger, Sonja Bisch-Knaden, Bill S. Hansson, Markus Knaden

## Abstract

Olfactory coding, from insects to humans, is canonically considered to involve considerable across-fiber coding already at the peripheral level, thereby allowing recognition of very large numbers of odor compounds. Here we show that the migratory locust, *Locusta migratoria*, has evolved an alternative strategy built on highly specific odorant receptors feeding into a complex primary processing center in the brain.

By collecting odors from food and different life stages of the locust, we identified 208 ecologically relevant odorants, which we used to deorphanize 48 locust olfactory receptors via ectopic expression in *Drosophila*. Contrary to the often broadly tuned olfactory receptors of other insects, almost all locust receptors were found to be narrowly tuned to one ligand and most of these best ligands exclusively excited only one receptor. Knocking out a single receptor using CRISPR-Cas9 resulted in an abolished physiological and behavioral response to the corresponding ligand. We conclude that the locust olfactory system, with an apparent lack of redundancy among most olfactory receptors, differs from so-far described olfactory systems.

## Introduction

The migratory locust *Locusta migratoria* is considered one of the world’s most harmful insects, whose swarms can devastate huge areas of crop and pasture (Uvarov, 1966; Zhang et al., 2019). The animals appear in two different phases –solitary and gregarious – that differ in morphological, physiological as well as behavioral traits, with only the gregarious phase forming swarms and causing agricultural problems (Wang and Kang, 2014; Zhang *et al*., 2019). Numerous behavioral and ecological laboratory and field studies have tried to decipher the phase shift from solitary to gregarious animals and by that targeted the formation of locust swarms and their biological control (Deng et al., 1996; Heifetz et al., 1996; Roessingh et al., 1998). Phase shift seems to be governed by population density, which in turn is affected by external factors like weather and food abundance (Cisse et al., 2013; White, 1976). In addition, phase shift is accompanied by up- and down-regulation of phase-specific genes (Wang and Kang, 2014) and seems to be triggered by intraspecific communication via visual, tactile, and olfactory cues (Ellis and Pearce, 1962; Gillett, 1968; Simpson et al., 2001). A well-investigated olfactory cue, turning locusts into the gregarious phase, is the so-called locustol (2-Methoxy-5-ethylphenol). This compound can be found in locust feces, and long-term exposure to it triggers the development of gregarious-specific behavioral and morphological traits in solitary locusts (Nolte et al., 1973). Furthermore, several aggregation pheromones like veratrole, guaiacol, and phenol that seem to be involved in the formation of swarms have been described (Fuzeau-Braesch et al., 1988). Another well investigated compound, phenylacetonitrile (PAN) has been suggested to both govern aggregation (Niassy et al., 1999) and repel predators (Wei et al., 2019). However, when revisiting the potential function of 35 compounds as aggregation pheromones (Guo et al., 2020), that before were identified in extracts either from locusts or their feces (Wei et al., 2017), only 4-vinylanisole turned out to carry this function for all locust developmental stages and both phases.

Olfactory cues, however, are not only involved in phase shift and aggregation, but, like in other insects, probably govern food- and reproduction-related behavior as well. During the process of maturation, volatile compounds emitted by male locusts have e.g. been shown to promote and synchronize the sexual development of both sexes, while compounds produced by young nymphs may retard the maturation of adults (Norris and Pener, 1965; Norris, 1962; Norris and Richards, 1964). Furthermore, volatile cues of solitary females have been shown to attract solitary males (Inayatullah et al., 1994), while volatiles in the so-called egg pods attract ovipositing females, resulting in a clustered distribution of eggs in the field (Saini et al., 1995). Finally, as migratory locusts mainly feed on gramineous plants (Despres et al., 2007; Wang et al., 2014), volatiles might help them to identify suitable host plants.

To detect and process all this olfactory information, locusts make use of an olfactory system that differs from those of other insects considerably. While the peripheral detection of odors via olfactory sensory neurons (OSNs) housed in basiconic, trichoid and coeloconic sensilla resembles that of other insects (Ochieng′ and Hansson, 1999; Yang et al., 2012), the first olfactory processing center, i.e. the antennal lobe, is very different (Ignell et al., 1998; Ignell et al., 2001). As in most animals, including humans, the insect antennal lobe contains spherical neuropil subunites, so-called glomeruli, where each glomerulus is usually targeted by OSNs expressing the same odorant receptor (OR) or olfactory ionotropic receptor (IR). From that, the number of glomeruli corresponds well with the number of ORs and IRs present in an insect. Hence, the identity of an odor is typically coded by the combinatorial activation of those glomeruli, whose receptors interact with the odorant. Food odorants often interact with many receptors, resulting in several activated glomeruli, while a few odors of specific ecological relevance become detected only by highly specific receptors and, hence, activate only single glomeruli (Haverkamp et al., 2018). In locusts, however, where 142 potential OR genes and 32 IRs have been annotated (Wang et al., 2015), the antennal lobe contains more than 1000 microglomeruli (Ignell *et al*., 1998; Ignell *et al*., 2001). This discrepancy between the numbers of olfactory receptors and glomeruli is a result of individual OSNs branching into several glomeruli (Hansson and Stensmyr, 2011; Ignell *et al*., 2001), a trait not observed in other insects. By allowing a much more diverse interaction between OSNs and projection neurons (PNs), the coding capacity of the locust could potentially be increased. However, the functional significance of such a system evolving from a glomerular architecture with unbranched OSNs and with most PNs targeting single glomeruli(Ignell *et al*., 1998; Ignell *et al*., 2001), into a system with thousands of microglomeruli innervated by highly branched OSNs and PNs is still unclear.

Here, we deorphanized 48 odorant receptors (LmigORs) that are available in full sequence and, hence, could be ectopically expressed in the *D. melanogaster* empty neuron system. In recordings from single sensilla (SSR), we functionally characterized these receptors using 208 odorants identified in the headspace of 4-5^th^ instar larval stages (from now on called nymphs), unmated and mated adults (both from the solitary and the gregarious phase) as well as from host- and nonhost plants, and thus of potential significance for *L. migratoria*. Surprisingly, contrary to other insects, where most receptors are broadly tuned (Carey et al., 2010; Guo et al., 2021; Hallem and Carlson, 2006; Pask et al., 2017; Slone et al., 2017; Wang et al., 2010), almost all investigated locust receptors were narrowly tuned to a single ligand and most of these best ligands activated only a single receptor. For several developmental stages and both sexes of gregarious and solitary locusts, we then tested the behavioral valence of 22 of such best ligands. We found that more than half of these compounds evoked significantly attractive or aversive behavior. More interestingly, the behavior evoked by an odorant strongly depended on the developmental stage, sex and/or phase of the tested animal. Finally, we used CRISPR/Cas9 to knock out LmigOR5, a receptor narrowly tuned to the aversive odorant geranyl acetone, a common locust- and plant-produced odorant. Animals lacking a functional LmigOR5 lost their antennal sensitivity to geranyl acetone and did not avoid it anymore.

From our results, we conclude that the unorthodox architecture of the locust antennal lobe is paralleled by a highly non-redundant receptor-ligand interaction, an architecture probably underlying a new type of olfactory information processing at these primary levels.

## Results and Discussion

### Odorants from ecologically relevant sources

Using solid phase microextraction-coupled gas chromatography-mass spectrometry (SPME-GC-MS) we identified volatile organic compounds from body and feces of all developmental stages and both sexes of both solitary and gregarious locusts as well as from two host- and two non-host plants, resulting in 32 samples (Fig. 1A+B). By comparing retention times and MS-spectra with those of synthetic standards, we were able to identify 186 compounds (of which 166 could be confirmed by their MS spectra) of potential ecological relevance (Fig. 1C). The compounds identified included the majority of those earlier reported from *L. migratoria* emanations, as e.g. PAN, 4-vinylanisole, guaiacol, benzaldehyde, phenol and 2,3-butanediol (Wei *et al*., 2017).

**Figure 1.**
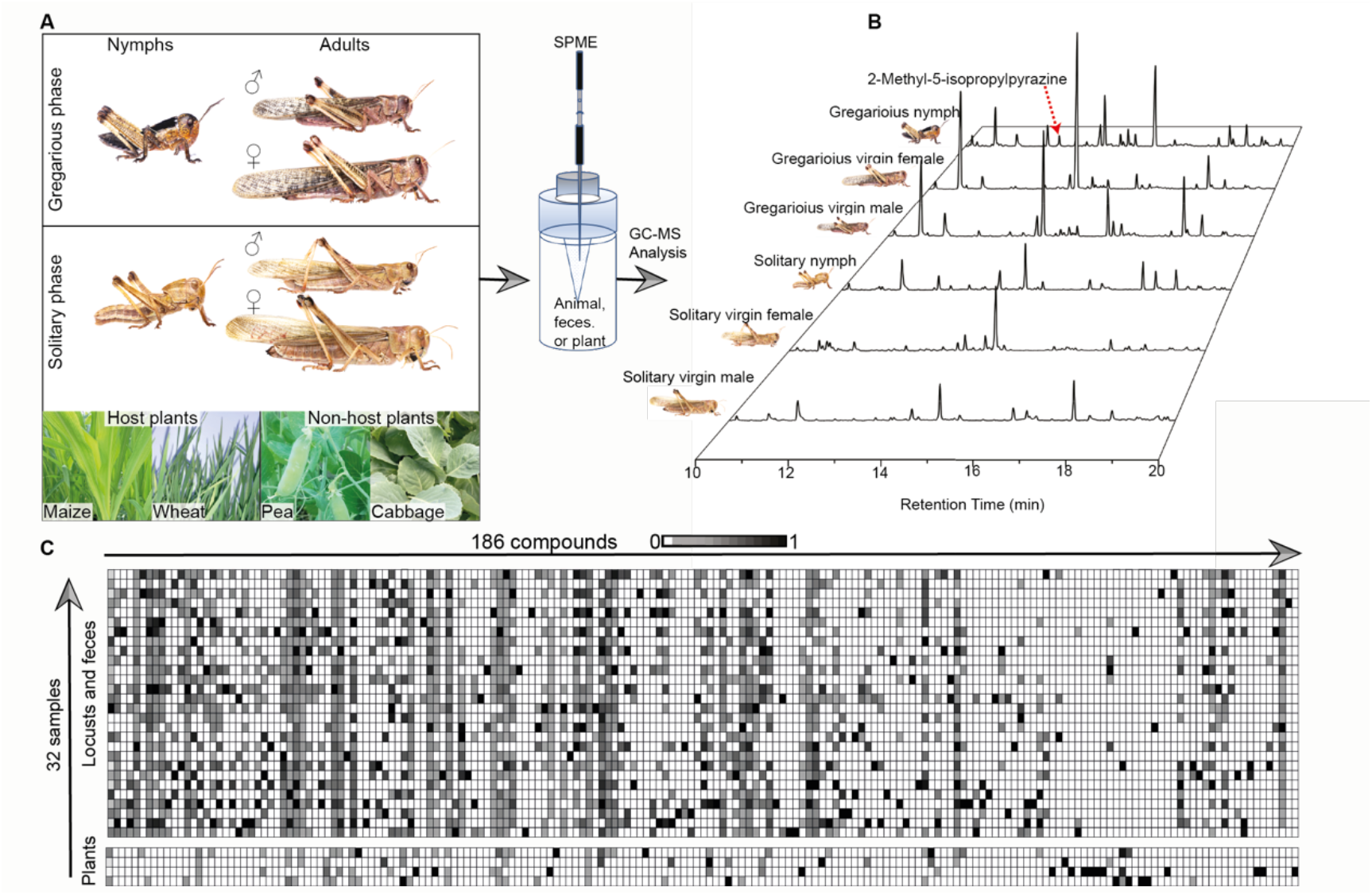
Odorants of potential ecological relevance. A. Example sources of which headspace samples were analyzed (i.e. 32 headspaces of both phases of nymphs, of immature, virgin, and mated females and males, and the corresponding feces of all of them, as well as 2 host- and 2 non-host plants). B. Gas chromatograms revealing 2-methyl-5-isopropylpyrazine as an odorant specific for 4-5 instar nymphs. C. Heat map presentation of 186 volatile compounds identified in the headspace of different sources normalized for each compound (N=4-8 per source; for a heat map presentation of additional, less volatile compounds that were identified in the bodies of nymphs and adults, see Suppl. Figure 1; for the quantification of all compounds see Suppl. Table 1).

In order to identify potential additional compounds of lower volatility, like cuticular hydrocarbons from locust body tissues, we used a thermal-desorption-unit coupled to GC-MS (TDU-GC-MS). We were able to identify 22 additional compounds (of which 21 could be confirmed by their MS spectra) from four body parts (hind leg, wing, abdomen and tergum) of the different developmental stages and both sexes of both solitary and gregarious animals (Suppl. Fig. 1). Taken together, we identified 208 compounds of potential ecological relevance (Suppl. Table 1).

When comparing the odorants emitted by virgin males and females of both phases, we could confirm an older study on stage-, sex-, and phase-specific odor emissions (Wei *et al*., 2017). Interestingly, we found additional novel odorants that were specific either for the sex (L-α-terpineol, emitted by gregarious and solitary males) or the mating state (2,4,6-trimethylpyridine, emitted by mated but not virgin gregarious females; E-2-octen-1-ol, emitted by virgin but not mated solitary females) (for a list of the abundance of odorants in different samples see Suppl. Table 1).

### Most *Locusta* odorant receptors are narrowly tuned

To identify *L. migratoria* odorant receptors (LmigORs) involved in the detection of the compounds identified, we next amplified the coding regions of 48 OR genes from antennal cDNA of adult locusts (phylogenetically covering the full range of annotated odorant receptor genes (Fig. 2A). The LmigOR genes were ectopically expressed in the *Drosophila melanogaster* empty neuron system (Fig. 2B). Previous in-situ hybridization studies in the desert locust *Schistocerca gregaria* had revealed that many OR genes are co-expressed with the sensory neuron membrane protein 1 (SNMP1) (Lemke et al., 2020). We, therefore, mis-expressed the receptors of interest in the *D. melanogaster* at1 empty neuron that is known to express SNMP (Kurtovic et al., 2007; Syed et al., 2010). From 48 cloned LmigORs, 43 turned out to be functional in the *Drosophila* at1 neuron, conferring a characteristic, regular, spontaneous firing rate (Fig. 2C). The remaining five receptors were non-functional as the at1 neurons displayed an abnormal spontaneous firing rate with bursts of action potentials, a phenotype reminiscent of that observed in mutant at1 neurons lacking its native receptor (Kurtovic *et al*., 2007). Next, we systematically examined the odorant detection spectrum of the 43 functional LmigORs by using the set of 208 locust- or food-derived compounds. At 10^−1^ dilution (i.e. 1 µL of a given compound dissolved in 10 µL solvent), all but one (LmigOR2) of the functional receptors exhibited significant responses (we used a cut-off of more than 15 spikes/s net increase) to at least one ligand. Of the 42 responding ORs, only one (LmigOR20) exhibited an extremely broad response pattern (Fig. 2C, for detailed response patterns of 42 receptors to 208 odorants see Suppl. Table 2), i.e. responded to more than 100 out of the tested 208 odors of different chemical classes. However, surprisingly, 41 LmigORs (98%) turned out to be narrowly tuned to a single or very few compounds (Fig. 2D, Suppl. Table 2).

**Fig. 2.**
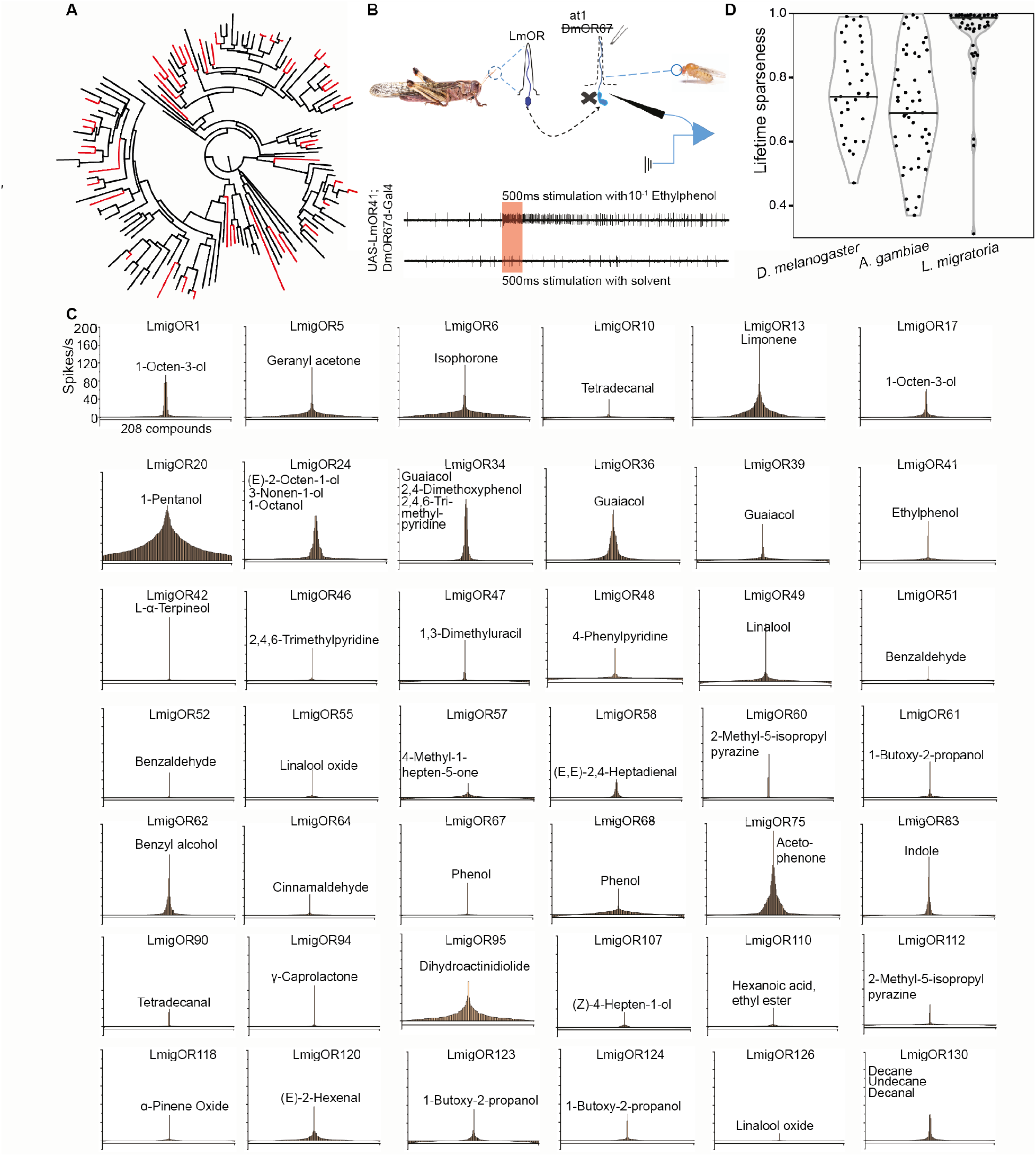
Response profiles of locust odorant receptors. A. Maximum likelihood tree based on amino acid sequences of 139 annotated odorant receptors genes of *L. migratoria*. B. Schematic of the heterologous expression system using an empty *Drosophila* at1 neuron with example traces from an at1 neuron mis-expressing LmigOR41 stimulated with ethylphenol (upper trace) or solvent (red bar, stimulus duration). C.Tuning curves of 42 receptors when stimulated with 208 compounds (tuning curves depict average responses of N=6-8 sensillum recordings per receptor). Compounds are displayed along the x-axis according to the strength of the responses they elicit from each receptor. Compounds eliciting strongest responses placed near the center with weaker responses towards the edges of the distribution. The most active identified ligands are depicted (for dose response characteristics of each receptor and its best ligand see Suppl. Fig. 2). D. Life-time sparseness of receptors of the fly *D. melanogaster* (n=35 receptors), the mosquito *Anopheles gambiae* (n= 50 receptors) and the locust *L. migratoria* (n=42 receptors) (filled violin plot differs from open plots, p<0.01, Kruskal-Wallis test with Dunn’s multiple comparison test).

The tuning width of an olfactory receptor can be calculated as its life time sparseness (LTS) (Bhandawat et al., 2007; Perez-Orive et al., 2002; Willmore et al., 2011) with values ranging from 0 to 1, with 0 signifying widely tuned, generalist receptors, while sparsely tuned, specialized receptors result in an LTS close to 1. With a median LTS of 0.96, the locust system exhibits a significantly more specialized set of receptors than described for other insect species investigated so far (*D. melanogaster*, 0.75 (Grabe et al., 2016); *Anopheles gambiae*, 0.69 (Carey et al., 2010), Fig. 2D).

In parallel to the highly specific LmigORs, each best ligand also only activates single or very few receptors (Suppl. Fig. 3). This means that the locust olfactory system offers fewer opportunities for across fiber coding at the first neural level. Therefore, it seems to be less redundant than in other insects, where many receptors are broadly tuned to several, chemically diverse ligands, and where single ligands are detected by many receptors (de Fouchier et al., 2017; Grabe *et al*., 2016; Hallem and Carlson, 2006).

### Behavioral screen of identified ligands

Narrowly tuned receptors are often suggested to respond to odorants of high biological relevance (Haverkamp *et al*., 2018). Without aiming for a detailed behavioral characterization of each identified ligand, we screened 22 of the identified best ligands for their general valence in a vertical two-way olfactometer (Fig. 3A). In this assay, a freely moving animal can decide between two rectangular areas (28cm x 28cm), of which one area contains the test odor, while the control area does not (Fig. 3B). By measuring the time spent in both areas, we calculated an attraction index ((time at odor − time at control)/total time) that could range from −1 (strongly aversive) to +1 (strongly attractive). We performed this experiment with 4-5^th^-stage nymphs, and virgin females and males of both phases.

**Fig. 3.**
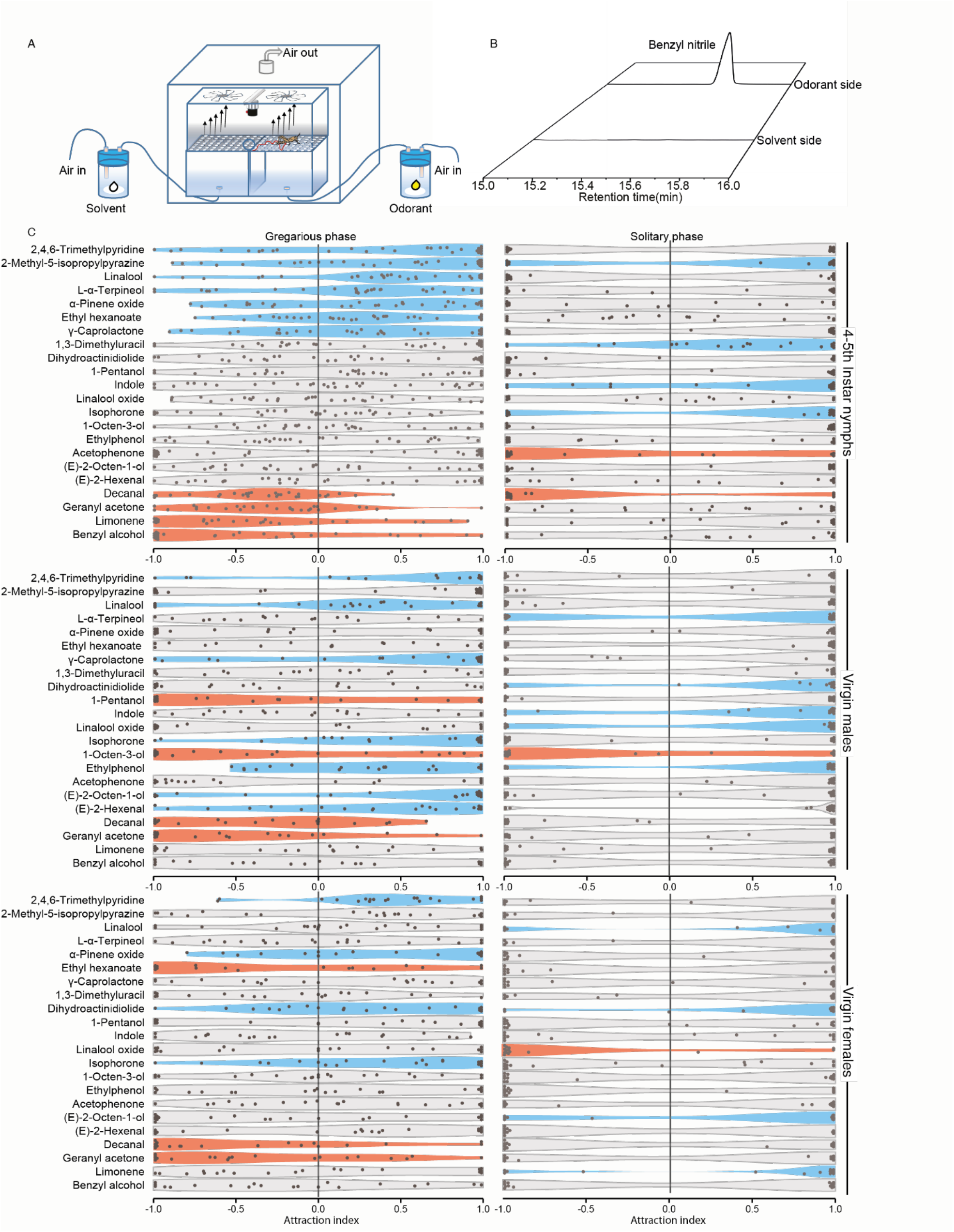
Behavioral responses of nymphs and virgin adults of both sexes to the identified best ligands. A. Vertical two-choice olfactometer, in which an animal can move within a solvent control side and an odorant-enriched test side. Test duration of individual animals, 10min. For details see Methods section. B. Solid-phase-micro-extraction followed by GC-analysis confirms odor-free control side and odor-enriched test side of the olfactometer. C. Attraction indices of gregarious (left) and solitary (right) nymphs (top), virgin males (middle) and virgin females (bottom) tested with the 22 identified best ligands. Attraction index = (time in odorant side − time in control side)/(time in odorant side + time in control side), with 1 signifying maximum attraction, and −1 signifying maximum aversion. Blue violins, significant attraction; red violins, significant aversion; Wilcoxon rank sum test against 0; gregarious nymphs, n=28-30 (tested individuals per odorant); gregarious males and females, n=20; solitary nymphs, males, and females, n=16.

Many of the tested odorants exhibited either attraction or repulsion to different subgroups of locusts (Fig. 3C). Geranyl acetone, an odorant emitted by all tested developmental stages and phases as well as by non-host plants (Suppl. Table 1) and detected by LmigOR5, repelled all gregarious stages, but was neutral to the solitary animals. We also found 2-methyl-5-isopropylpyrizine (an odorant characteristically emitted in high concentration from both gregarious nymphs and their feces (Suppl. Table 1) to be specifically attractive to the nymphs of both phases but neutral to all tested adults. However, the ecological relevance of these odorants and their attractiveness to specific subgroups of locusts remains elusive. Interestingly, while some of the compounds elicited strong responses in one or several subgroups of animals, none of them was attractive to all animals, as could have been expected from e.g. a odor like linalool or linalool oxide, that are strongly emitted by host plants.

Typical pheromone receptors of other insects have been shown to be as narrowly tuned as locust odorant receptors (Fleischer and Krieger, 2018; Kurtovic *et al*., 2007). We, therefore, hypothesized that some of the deorphanized LmigORs would be involved in pheromone detection. Consistent with this hypothesis, some receptors were narrowly tuned towards compounds that, based on their presence in a specific sex, phase, or developmental stage, potentially could fulfill a pheromonal role. LmigOR46, for example, responded narrowly to 2,4,6 trimethylpyridine, a compound emitted by gregarious females only after mating and, therefore, potentially involved in post-copulatory mate guarding, as described for several drosophilid flies (Khallaf et al., 2020; Meer et al., 1986). Furthermore, LmigOR24 was tuned to E-2-octen-1-ol (emitted by virgin but not mated solitary females) and LmigOR42 to L-α-terpineol, a male-specific odor in both the gregarious and the solitary phase (Suppl. Table 1).

None of the behavioral results, however, pointed at a pheromonal function of any of the tested odorants. The aforementioned 2,4,6-trimethyl pyridine for example, an odor emitted mainly by mated gregarious females (Fig. 1), but in lower amounts also by other subgroups and their feces (Suppl. Table 1) and detected by OSNs expressing LmigOR 46 (Fig. 2), was significantly attractive to all tested gregarious stages, while eliciting no response in any solitary stages (Fig. 3C). Furthermore, E-2-octen-1-ol (emitted by virgin but not mated solitary females) or L-α-terpineol (emitted only by males of both phases) (Suppl. Table 1) did not elicit a specific response in the opposite sex (Fig. 3C), suggesting that these odorants either do not play a major role in sexual behavior or do so only in the context of other cues.

Another characteristic trait of sex pheromone receptors is that their expression often is upregulated or even specific for the receiving sex (Bray and Amrein, 2003). We, therefore, analyzed the expression patterns of ORs in different developmental stages and both sexes of both the solitary and the gregarious phase to investigate whether any of the aforementioned LmigORs that were tuned to sex- or mating state-specific odorants would be upregulated in the potential recipient of this information. To do so, we examined the mRNA expression levels of olfactory receptors using NanoString nCounter, a technology primarily developed for gene expression analysis (Goytain and Ng, 2020) (for a detailed list of expression patterns of ORs and IRs in gregarious and solitary nymphs, virgin females, and virgin males see Suppl. Table 3).

As the expression level of locust receptors detecting 2,4,6 trimethylpyridine (LmigOR34), E-2-octen-1-ol (LmigOR24), and L-α-Terpineol (LmigOR42) was not differential for a given developmental stage or sex (Suppl. Table 1) it does not support any specific involvement in pheromone interactions. However, behavioral responses to a given pheromone do not necessarily rely on sex-, stage- or phase-specific expression of pheromone receptors, but can also be due to processing differences at higher brain centers (Ruta et al., 2010). Therefore, our analysis does not rule out the existence of pheromones in *L. migratoria. Drosophila* larvae (Fishilevich and Vosshall, 2005) and adults (Elmore et al., 2003; Trimmer et al., 2019) exhibit behavioral attraction to a given food odorant, even when lacking an olfactory receptor detecting this odorant. The reason behind these results seems to be the redundancy of the *Drosophila* system, where olfactory coding is based on the participation of many receptors that detect partly overlapping sets of odorants, so-called across-fiber coding (Hallem and Carlson, 2006). Therefore, receptors with overlapping tuning profiles can compensate for the absence of a single receptor. Having shown that in the locust most LmigORs are narrowly tuned (Fig. 2C) and that many odorants seem to be detected by single or few ORs, we next aimed to probe the presence or lack of redundancy in locust odorant detection. As geranyl acetone provoked strong repellency in gregarious locusts (Fig. 3), we asked whether knocking out LmigOR5, a receptor specifically tuned to geranyl acetone, would harm the locust detection of and behavior towards this odorant. To address this question, we generated an LmigOR5 mutant line using CRISPR-Cas9 genome editing (Fig. 4A). The obtained LmigOR5^-/-^ mutant line contained a 149-bp deletion with a 59-bp deletion in non coding 5’ UTR part and a 90 bp deletion in the exon1. As the remaining part of the gene has a start codon 18 bp downstream of the deletion, the deletion results in a protein lacking in total 36 amino acids. When recording either from the whole antenna (electroantennogram recordings, EAGs), or from basiconic sensilla (single sensillum recordings, SSRs), mutant locusts detected odorants not related to LmigOR5 (isophorone and 2,4,6-trimethylpyridine) like wild type animals (Fig. 4B,C). They also exhibited strong attraction towards 2,4,6 trimethylpyridine, indicating the lack of any relevant phenotypical off-target effects of the CRISPR-Cas9 manipulations (Fig. 4D). However, the electrophysiological response to geranyl acetone, both at the level of the antenna and the sensillum (Fig. 4B-C), as well as the wild-type-typical avoidance of this odor (Fig. 4D) was fully abolished in the mutant animals, suggesting the necessity of LmigOR5 for the detection of geranyl acetone and the corresponding behavior. No across-fiber coding patterns did thus ameliorate the effects of the missing receptor.

**Fig. 4.**
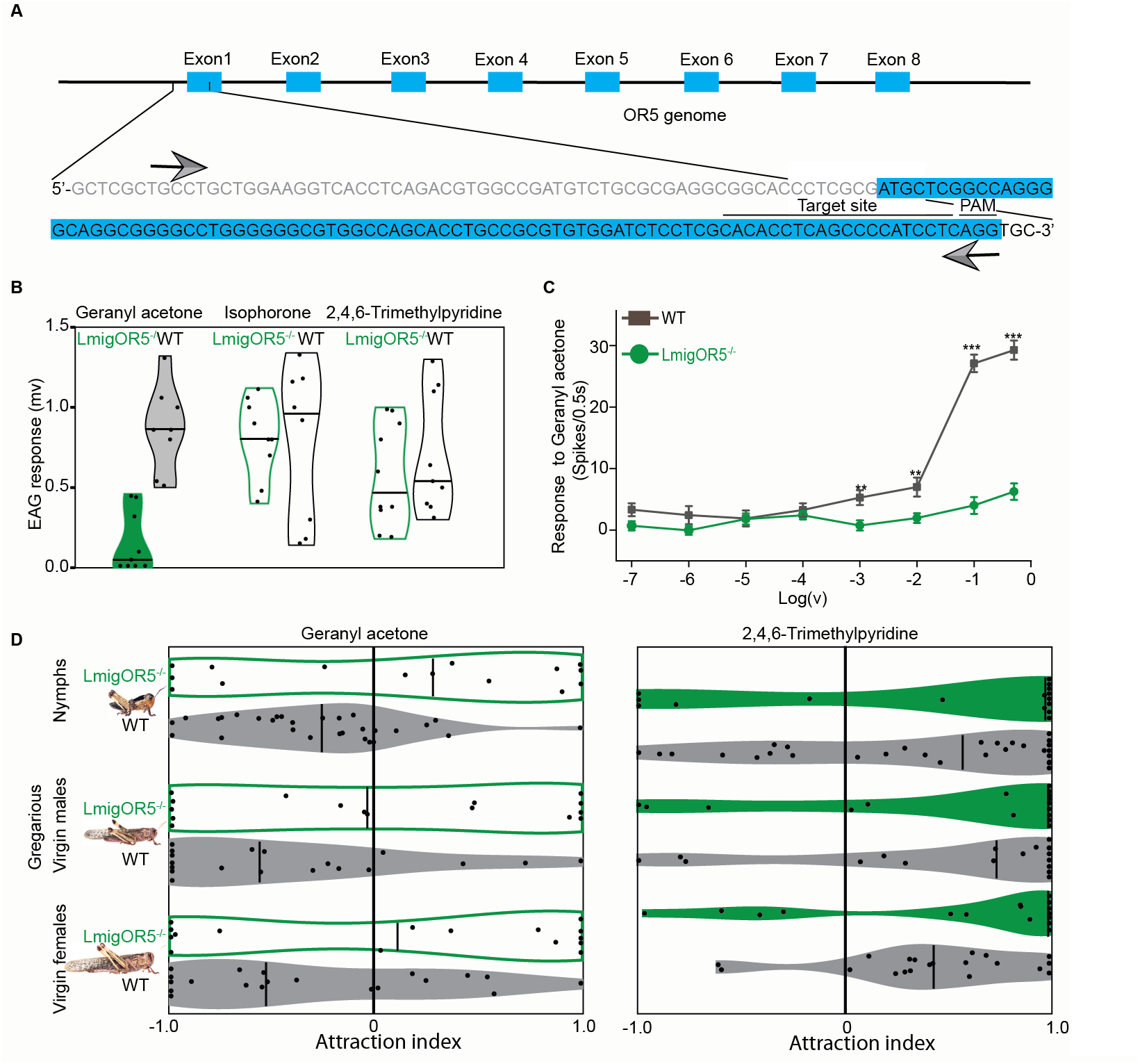
LmigOR5 is necessary for detection of and aversion to geranyl acetone. A. Schematic of the CRISPR-Cas9 strategy to knock out LmigOR5. Exons 1-8, coding regions of the LmigOR5 gene; nucleotide sequence depicts the CRISPR-Cas9-induced deletion of 149bp of the non-coding (grey letters) and coding (blue-shaded letters) region; deleted part depicted by arrows; target site and the proto adjacent spacer motive (PAM) are highlighted. B. Electroantennographic (EAG) responses of LmigOR5-knockout animals and wildtype animals to the LmigOR5 ligand geranyl acetone and two control odorants (filled boxes differ from each other (p<0.01,N=8-10 per test; Mann Whitney U-test; black lines, median). For EAG dose-response curves see Suppl. Fig. 4. C. Single sensillum recording (SSR) dose response curves for geranyl acetone in LmigOR5-knockout and wildtype locusts. SSR experiments were performed from basiconic sensilla (knock-out animals, n=20 recordings; wildtype animals, n=19 recordings; dots, mean; error bars, standard error; Mann Whitney U-test). D. Left panel, lack of repulsion of geranyl acetone towards LmigOR5 knockouts (green violins) as compared to wildtype animals (grey violins). Right panel, wildtype-like behavior of LmigOR5 knockouts towards the attractant 2,4,6-trimethylpyridine, detected by LmigOR46. All experiments performed with gregarious animals. Filled violins, significant attraction or repulsion (Wilcoxon rank sum test, knock-out animals, n=15-16, wild-type animals, n=20-30, filled violins significantly differ from 0).

We conclude that as most of the 42 deorphanized LmigORs, which well represent the genetic diversity of all 142 annotated odorant receptor genes (Fig. 2A), were narrowly tuned to very few compounds, the olfactory sensory system of *L. migratoria* exhibits a very low redundancy and by that might differ dramatically from other so-far described olfactory systems. Up to now, it was suggested that insects as well as vertebrates can identify a virtually infinite number of odors by using a combinatorial code, in which one OR recognizes multiple odorants and one odorant is recognized by multiple ORs, so that different odorants are recognized by different combinations of ORs (Malnic et al., 1999). As such a strategy demands numerous broadly-tuned odorant receptors, and as only one out of 43 deorphanized LmigORs (LmigOR20 responding to >100 out 208 odorants) fulfilled this demand, *L. migratoria* obviously has evolved an alternative way to code odor information. One important role might play temporal coding, i.e. information about odor identity might not only be coded by the combination of responding OSNs, but also by the temporal characteristics of their individual responses (Laurent, 1999; Laurent et al., 2001). Although the final strategy of olfactory coding in locusts remains elusive, it might very well relate to the enormously complex structure of the locust antennal lobe (Ignell *et al*., 1998; Ignell *et al*., 2001), where highly complicated connections between neurons might allow the equivalent resolution to be gleaned as in a system built on first order across-fiber coding. Future investigations will reveal whether locusts can discriminate different odorants as well as insects having access to a redundant system and by that a combinatorial code at their disposal.

## Methods

### Insects

The gregarious and solitary locusts (*Locusta migratoria*) used in the experiments were reared at the Max Planck Institute for Chemical Ecology. In brief, gregarious locusts were reared in cages (30 cm × 30 cm × 30 cm) with 400 to 500 first-instar locusts per cage in a well-ventilated room. Solitary locusts were raised in single cylindrical box (10.5cm high × 8cm diameter), each box with a separate ventilation system. Both the gregarious and solitary locusts were maintained for at least three generations before the experiments were conducted. All locusts were cultured under the following conditions: a 14 h:10 h light:dark photoperiod, temperature of 30 ± 2 °C, relative humidity of 50 ± 5%, and a diet of fresh, greenhouse-grown wheat seedlings for gregarious locusts and solitary locusts.

### Chemical analysis

The volatile compounds from all these different samples were collected by solid phase microextraction (SPME) for 10 hours at 30 °C. To compare volatile profile of gregarious and solitary locusts in nymphs (fourth to fifth instar) and adult stages (immature adult refers to 2-day-old post-adult eclosion, mature adult is 10-day-old post-adult eclosion before mating, mated adult was 1-day-old after mating), a small number of locusts (6 individuals for nymphs or 2 individuals for adults) were confined into a 100 ml glass bottle, a SPME fiber (PDMS/DVB 65 *µ*m) was introduced inside of bottle through the cap and a PEAK guide piece which served as barrier to avoid locust direct contact with fiber. Meanwhile, 300mg fresh feces were collected from each stage of locusts and enclosed in a 1.5 ml Agilent vial, the fiber was exposed to head space by directly penetrating the septum of the cap with the SPME fiber holder. Plant materials were subjected to the similar SPME system that applied to body volatile, approximately 5g fresh leaves were collected from 4 plant species including two host plants (*Zea mays* and *Triticum aestivum*) and two non-host plant (*Brassica oleracea* and *Pisum sativum)* and placed individually in 500ml glass bottle for subsequent fiber collection. For each type of odor collection, the SPME volatiles collected from an empty glass vial for 10 hours served as control. After each odor collection, the SPME fiber was retracted and immediately inserted into the inlet of a gas chromatography–mass spectrometry (GC–MS) system equipped with an HP5 column. After fiber insertion, the column temperature was maintained at 40 °C for 3 min and then increased to 150 °C at 5 °C·min^-1^, thereafter it was increased to 260 °C at 10 °C·min^-1^, followed by a final stage of 5 min at 260 °C. Compounds were identified by comparing mass spectra against synthetic standards and NIST 2.0 library matches. Most of the synthetic odorants that were tested and confirmed were purchased from Sigma–Aldrich.

### Cloning of Locust ORs

To clone as much as possible full-length coding sequences of locust ORs, we collected > 50 antennae and palps from 7-day-old post-adult in both G and S phase. RNA of each sample was extracted using an RNA isolation kit (QIAGEN) according to the manufacture’s instructions. First-strand complementary DNA (cDNA) was generated from 5 µg of total RNA pooled with equivalent amount of all types of olfactory tissue samples using oligo dT-primed cDNA synthesis with Superscript III Reverse Transcriptase (Invitrogen) for the generation of templates for subsequent PCR reactions. PCR was performed with MyTaq DNA or Myfi Polymerases (Bioline) and primers were designed (Suppl. Table 4) according to published sequences available at GenBank^25^. PCR amplification products were separated on a 1.0% agarose gel and were cloned into the PCR 2.1 TA cloning vector (Invitrogen) and verified by sequencing. Sequences of at least 4 independent clones were obtained for each OR and compared to verify polymorphisms as such rather than PCR errors. The cloning OR sequences which have been used for functional analysis are listed in supplementary material.

### Transgenic expression of LmigORs in *D. melanogaster* at1 neurons

The entire coding region of each LmigOR was sub-cloned into an empty neuron vector pUAST.attb (a gift from J. Bischof) using two combinations of restriction enzymes kpnI/XbaI or EcoRI/XhoI (New England Biolabs). Homozygous UAS-ORX lines (with transgene insertions into chromosome II) were generated at Bestgene (https://www.thebestgene.com). An OR67d-GAL4 stock (provided by B. J. Dickson) was individually crossed to each of the transgenic UAS-Lm-ORX flies, and homozygous lines expressing the Or gene of interest in the decoder at1 neuron of *D. melanogaster* were established. Each UAS-transgenic line was confirmed by sequencing of genomic DNA prepared from the final stocks.

### Single-Sensillum Recordings (SSR)

To test the function of individual locust odorant receptors in the *Drosophila* empty neuron system, we performed SSR recordings from fly at1 sensilla according to standard procedures(Khallaf *et al*., 2020). Briefly, adult flies were immobilized in 200ul pipette tips, and the third antennal segment was placed in a stable position onto a glass coverslip. The at1 sensillum type was identified under a microscope (BX51WI; Olympus) at ×100 magnification and tested with odors listed in Suppl. Table 2. The extracellular signals originating from the OSNs were measured by inserting a tungsten wire electrode into the base of a sensillum and a reference electrode into the eye. Signals were amplified (Syntech Universal AC/DC Probe; Syntech), sampled (96000 samples/s), and filtered (500 to 3000 Hz with 50/60-Hz suppression) via USB-Universal Serial Bus-Intelligent Data Acquisition Controller (IDAC) (Syntech) connected to a computer. Action potentials were extracted using AutoSpike software, version 3.7 (Syntech). Synthetic compounds were diluted in dichloromethane, hexane or mineral oil (Sigma-Aldrich, Steinheim, Germany). Before each experiment, 10 µl of diluted odor was freshly loaded onto a small piece of filter paper (1 cm^2^, Whatman, Dassel, Germany) and placed inside a glass Pasteur pipette. The odorant was delivered by inserting the tip of the pipette into a constant, humidified airstream flowing at 600ml/min through a 8mm inner diameter stainless steel tube ending 1cm from the antenna. Neural activity was recorded for 10 s, starting 3 s before the stimulation period of 0.5 s. Responses from individual OSNs were calculated as the increase (or decrease) in action potential frequency (spikes per second) relative to the pre-stimulus frequency. Traces were processed by sorting spike amplitudes in AutoSpike, analysed in Excel, and illustrated using Adobe Illustrator CS (Adobe systems, San Jose, CA). In addition, we conducted SSRs from locust basiconic sensilla with functional ligands identified in the *Drosophila* empty neuron recording. Each locust was confined in a plastic tube 1ml in diameter and its antennae were fixed with dental wax. The recording and analysis process underwent the same treatment as in the fly.

### Electroantennograms (EAGs)

Electroantennograms (EAGs) of OR5 mutant locusts and wildtype animals were performed by cutting off the antenna of male animals the bases of the flagellum. The cut end was immediately placed into a glass capillary containing locust ringer solution (140 mM NaCl, 5 mM CaCl_2_, 5 mM KCl, 4 mM NaHCO_3,_ 1 mM MgCl_2_, 6.3 mM Hepes, 15 mM Sucrose). Both capillaries were then placed on Ag-AgCl wires of which one was connected to a grounding electrode, while the other was connected to a 10x a high-impedance d.c. amplifier. Both capillaries were then placed on Ag-AgCl wires, of which one was connected to a grounding electrode, while the other was connected to a 10x a high-impedance DC amplifier (Syntech; The Netherlands). The signal was then sent to an analog/digital converter (IDAC-4, USB, Syntech) and transferred to a computer. Finally, the data were analyzed and saved by using AutoSpike software, version 3.7 (Syntech). The odorant was delivered by inserting the tip of the pipette into a constant, humidified airstream flowing at 600 ml/min through an 8 mm inner diameter stainless steel tube ending 1cm from the antenna. The response was recorded for 10 s, starting 3 s before the stimulation period of 0.5 s.

### Locust behavioral responses and video-tracking system

Dual-choice olfactometer experiments were conducted as shown in Figure 4A. We used a vertical airflow olfactometer, similar to the architecture described in a previous study (Wei *et al*., 2019). Two plastic containers (28 cm × 28 cm × 18 cm) with open top, seamlessly connected to a glass chamber (60 cm × 30 cm × 30 cm) constituted the main structure of the behavioral observation chamber. The top of each container was equipped with a plastic plate. These plates had small holes 1 mm in diameter at 1 cm distance from each other. The bottom of each container was connected to an air purification system consisting of a compressed air cylinder, a charcoal filter and a molecular sieve filter. A flowmeter guaranteed a constant rate of airflow (3l/min) through each plastic container at each side (zone) of the arena. The glass chamber enclosed the area above the two plates and thus formed the behavioral observation area. The top of the chamber was equipped with two fans to provide vertical airflow and a video camera was installed in the gap between the fans. The bioassay setup was placed in an observation room (60 cm × 70 cm × 80 cm) with a ventilation system at the top. White light panels were located in the ceiling to provide uniform lighting. The bioassay provided two choices for locusts tested: a clean, vertical airflow in the control zone and an adjacent vertical airflow filled with the odor tested. For the series of behavioral tests, locusts entered the arena through a small door in the middle of the Plexiglas chamber and were allowed to stay in the olfactometer for 10 min. The diluted odorant was applied to a piece of filter paper (3 cm × 3 cm; Whatman No. 1), and paraffin oil was applied in a similar way to serve as a control. After testing 8–15 individuals, the positions of odor and control were reversed to prevent position deviation. The container was then cleaned with 75% ethanol and ventilated for 30 min to remove any odor residues. By using a HD digital video camera, combined with media recorder software (Media recorder 2.5), we captured the locusts’ behavioral activities during 10 min at 30 frames s^−1^ after introduction into the arena. Video recordings were analyzed by manually observing the total time spent on each side (unit: s). Valence of the tested odorants was quantified with an attraction index (AI), calculated as: AI = (O-C)/(O+C), where O is the time of the locust spent in the odorant panel and C the time spent in the control panel.

### Gene expression analysis

Antennal tissues from stage 4 nymphs and virgin adult males and females of both solitary and gregarious phase *L. migratoria* were collected in 2 ml microcentrifuge tubes, immediately immersed in liquid N_2_, and stored at -80°C until further processing. Total RNA was extracted from the tissues using RNeasy Mini Kit (Qiagen, Germany) and RNase-free DNase Set (Qiagen, Germany) was used for the DNase digestion step. The concentration and purity of the extracted total RNA were checked on a NanoDrop™ One (Thermo Scientific) and three samples for each experimental group were selected.

We used the nCounter Elements XT gene expression assay (NanoString Technologies, Inc, USA). Probes A and B for the target genes were designed based on sequences published on GenBank. Off-targets were checked by BLAST to Locust genome assembly v2.4.1 predicted coding sequences (http://locustmine.org/index.html).Master stocks for Probe A pool and Probe B pool were generated by Integrated DNA Technologies (IDT, Inc.). TagSets for 192 targets and buffers were purchased from NanoString Technologies, Inc, USA. The hybridization reaction (15 µl) was set-up using standard protocol (MAN-10086-01, Page 16). Based on initial standardization, 300 ng total RNA was used. Hybridization was done at 67°C for 16 h, after which 20 µl Merck water was added to the sample. 30 µl of each sample was then loaded on nCounter SPRINT Cartridge (NanoString, USA) and processed on nCounter SPRINT Profiler (NanoString, USA).

The raw data from the gene expression assay was processed using nSolver4.0 (NanoString, USA). Quality control for mRNA data was done on each sample using default parameters for nCounter SPRINT Profiler according to NanoString Gene Expression Data Analysis Guidelines (MAN-C0011-04). The parameters were Imaging QC: 75; Binding Density QC: 0.1 – 1.8; Positive Control Linearity QC: 0.95; Positive Control Limit of Detection QC: 2 standard deviations. A background subtraction was done first using the raw counts of the 8 negative control probes (mean + 2 standard deviations). After that, two normalization steps were performed, first using the geometric mean counts of the 6 external positive control probes, and second, using the geometric mean counts of two endogenous reference genes (18S rRNA and EF1-alpha). The geometric mean normalized data are given in Supplementary Table S3.

### Establishment of the LmigOR5 mutant line using CRISPR–Cas9

The establishment of LmigOR5 mutant locusts by CRISPR–Cas9 was performed as previously described (Guo *et al*., 2020). In brief, the embryos of locusts were collected from egg pods, washed with 75% ethanol, and were placed on 1% agarose gel. The purified Cas9 protein and single guided RNA were mixed to final concentrations of 400 and 150 ng µl^−1^, respectively, and 27.2 nl were injected into the embryos using a nanolitre injector (World Precision Instruments) with a glass micropipette tip under an anatomical lens (for the target site see Fig. 4A). Then, the embryos were placed in a 30 °C incubator for approximately 14 days until the locusts hatched. The first-instar nymphs were placed in cages (30 cm × 30 cm × 30 cm) with a 14 h:10 h light:dark cycle and sufficient food. In order to screen for successful mutations, we collected parts of adult legs and lysed them with a 45 µl NAOH buffer (50 mM) at 95 °C in a PCR machine for 30 min and added 5 µl Tris-HCL (pH 8.0, 1 M). Then, we used a 2 µl template to amplify the targeted fragments and sequenced the fragments to identify whether the mutants were generated. The PCR reaction volume contained 5µL 5x MyTaq Reaction Buffer, 2µL (10 µM) F primer, 2ul (10 µM) R primer, 0.5 µL MyTaq HS DNA polymerase, 13.5µL nuclease free water and 2µL template. The PCR reaction condition was 95°C 1min, 40 cycles of 95°C 15s, 60°C 15s and 72°C 30s, followed by a final 10min extension period of 72°C. In 78 locust individuals, 16 locusts with mutations in exon 1 (mutation efficiency: 20.5%) were identified. To further investigate the exact mutation models, we performed Sanger sequencing of PCR amplicons from all mutated locust individuals. Mutations in G0 locusts were evaluated by using PCR-based genotyping (for primer information see Suppl. Table 4). G0 mutants were crossed with the wild type to obtain G1 offspring. G1 locusts, whose DNA strands contained a 149-bp deletion, were crossed with each other to establish stable lines. For the expanding mutant population, 149-bp-deleted homozygotes of G2 locusts were further crossed with each to generate a line of homozygotes G3 animals. Finally, OR5 homozygous mutant lines were established successfully.

### Polygenetic tree generation

To construct the LmigOR phylogenetic tree, a total of 139 amino acid sequences from *L. migratoria* were aligned with MAFFT. The phylogenetic tree was built with FastTree version 2.1.3 using approximated maximum likelihood method and the resulting tree was visualized using FigTree version 1.4.2.

### Statistics and figure preparation

All statistics were analyzed using GraphPad Instat (https://www.graphpad.com/scientific-software/instat)and preliminary figures were conducted using PAST (https://past.en.lo4d.com). Figures were then processed with Adobe Illustrator CS5.

## Author contribution

HC, BSH, and MK designed study and wrote the manuscript; HC and KW collected and analyzed volatiles; HC and SBu cloned and deorphanized odorant receptors; AU established behavioral assay; HC conducted behavioral experiments, HC, AU and SBr collected animal tissue and performed RNA extraction for gene expression analysis; MTT, SB-K, and LCL performed gene expression analysis.

## Acknowledgments

We thank Kang Le for helpful information on the <u>locust transcriptome and genomic data. The study was funded by the Max Planck Society</u>.

## Supplemental information

**Suppl. Figure 1.**
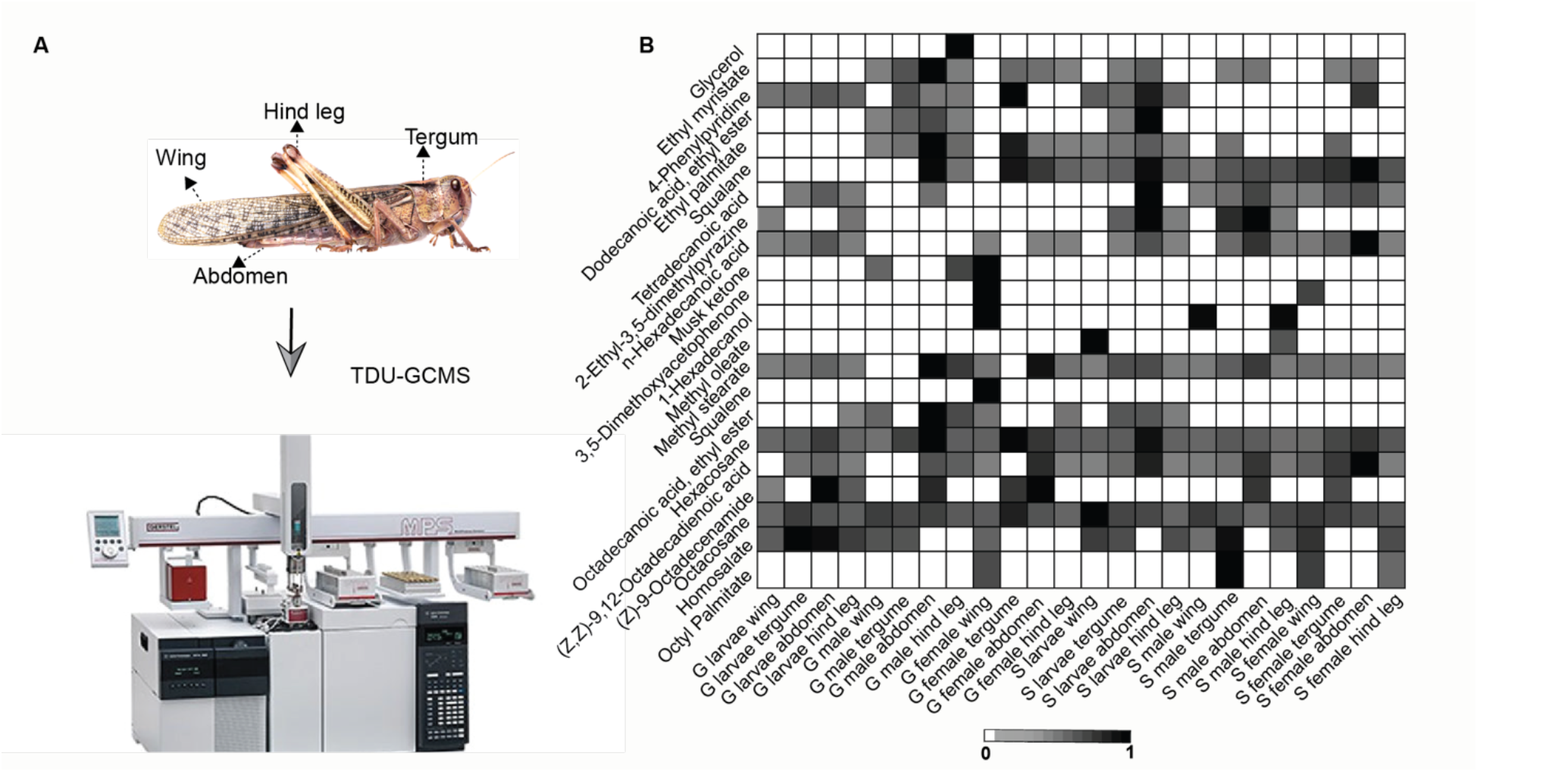
Cuticular hydrocarbons of potential ecological relevance. A. Example sources (highlighted body parts of different stages, phases, and sexes of locusts) that were analyzed in the thermal-desorption-coupled GC-MS. B. Heat map presentation of the 22 compounds identified in the different samples (N=4-8 per source).

**Suppl. Figure 2.**
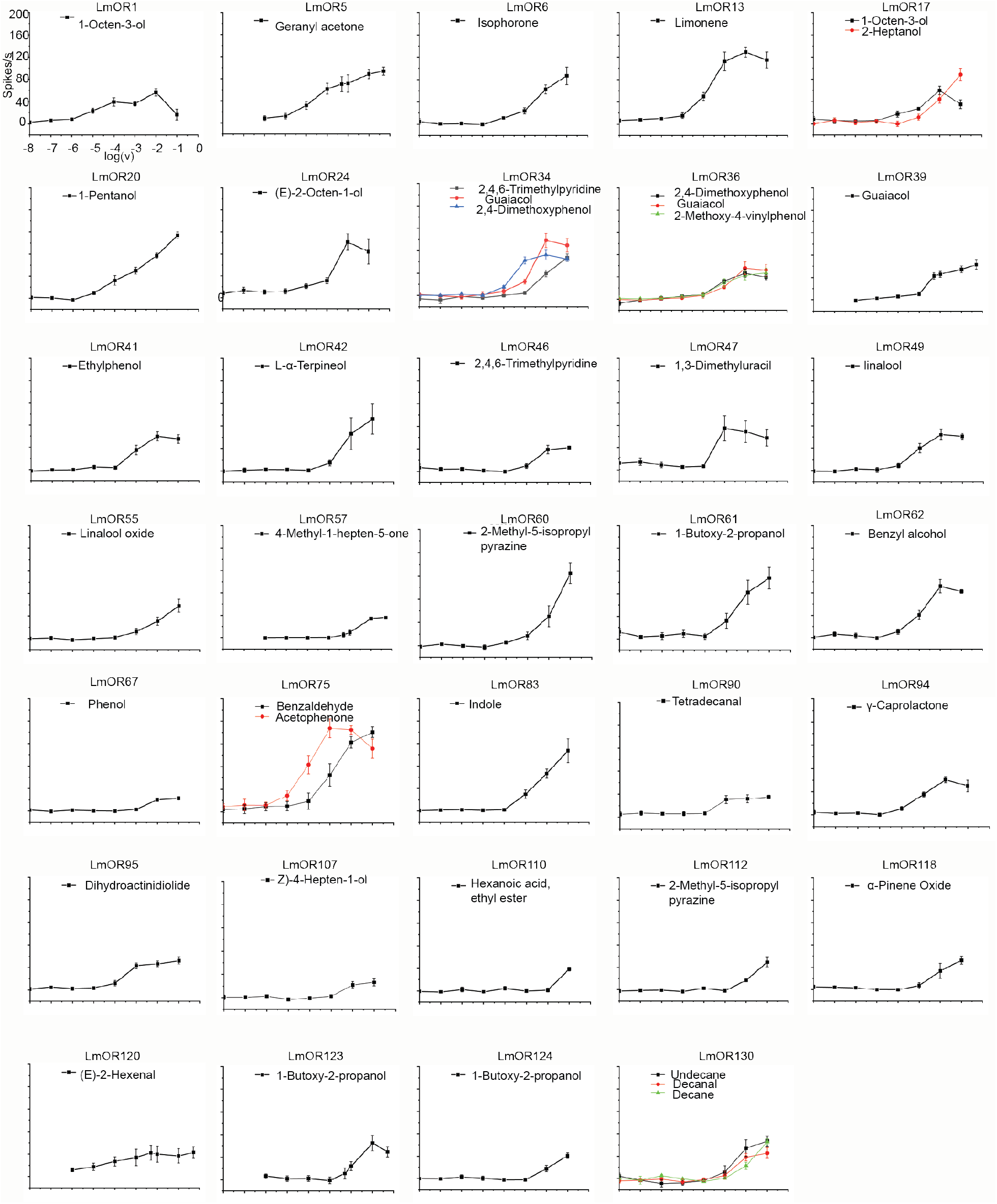
Dose response interactions of LmigORs with identified best ligands. Response curves depict average responses (and SEM) of N=4-7 sensillum recordings per receptor and ligand.

**Suppl. Figure 3.**
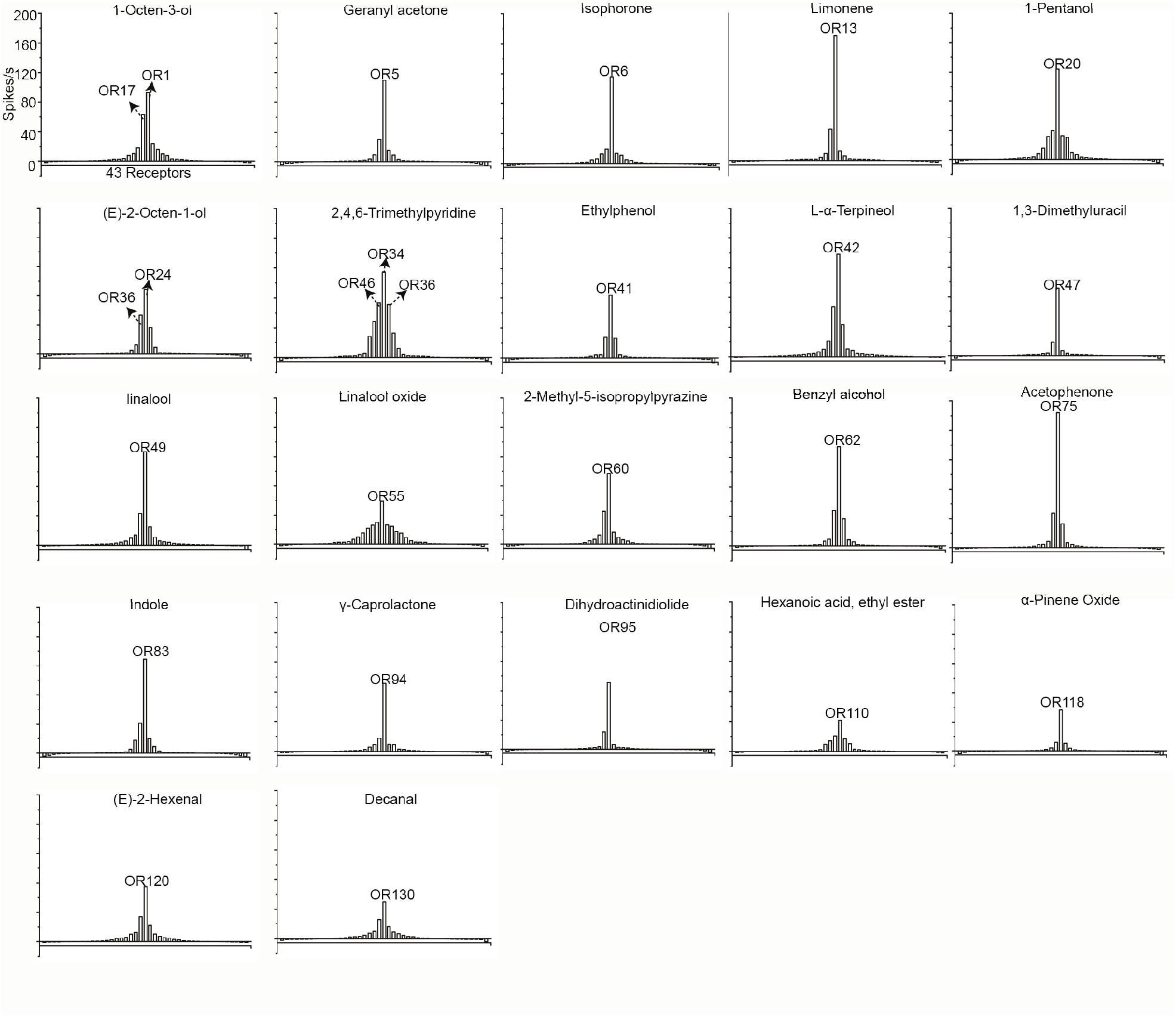
Tuning curves for odorants. The responses of the 43 locust ORs are ordered along the X-axis according to the magnitude of the response they generate for a given compound. Strongest (weakest) responding receptors placed near the center (edges) of the distribution. Strongest responding receptors are depicted.

**Suppl. Figure 4.**
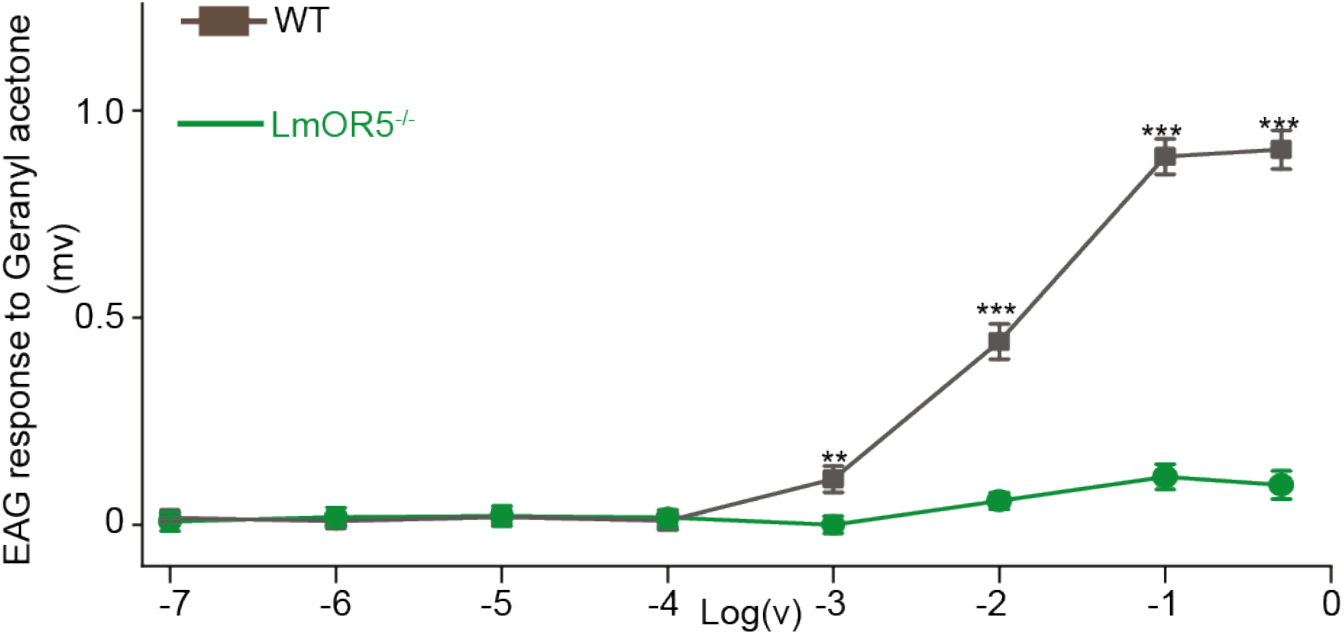
Electroantennogram dose response curves for antennal activation with geranyl acetone. N=15 (LmigOR5 mutants) and 12 (wildtype animals). Curve depict average and SEM. Grey, wildtype responses; green, responses of LmigOR5 knockout animals.

